# Migraine aura, a predictor of near-death experiences in a crowdsourced study

**DOI:** 10.1101/811885

**Authors:** Daniel Kondziella, Markus Harboe Olsen, Coline L. Lemale, Jens P. Dreier

## Abstract

**Background:** Near-death experiences (NDE) occur with imminent death and in situations of stress and danger but are poorly understood. Evidence suggests that NDE are associated with rapid eye movement (REM) sleep intrusion, a feature of narcolepsy. Previous studies further found REM abnormalities and an increased frequency of dream-enacting behavior in migraine patients, as well as an association between migraine with aura and narcolepsy. We therefore investigated if NDE are more common in people with migraine aura.

**Methods:** We recruited 1037 laypeople from 35 countries via a crowdsourcing platform. Reports were validated using the Greyson NDE Scale.

**Results:** Eighty-one of 1037 participants had NDE (7.8%; CI 6.3-9.7%). There were no significant associations between NDE and age (p>0.6, t-test independent samples) or gender (p>0.9, chi-square test). The only significant association was between NDE and migraine aura: Forty-eight (6.1%) of 783 subjects without migraine aura and 33 (13.0%) of 254 subjects with migraine aura had NDE (p<0.001, chi-square test, odds ratio (OR) = 2.29). In multiple logistic regression analysis, migraine aura remained significant after adjustment for age (p≤0.001, OR 2.31), gender (p<0.001, OR 2.33), or both (p<0.001, OR 2.33).

**Conclusions:** In our sample, migraine aura was a predictor of NDE. This indirectly supports the association between NDE and REM intrusion and might have implications for the understanding of NDE, because a variant of spreading depolarization (SD), terminal SD, occurs in humans at the end of life, while a short-lasting variant of SD is considered the pathophysiological correlate of migraine aura.

## Introduction

Near-death experiences (NDE) include emotional, self-related, spiritual and mystical perceptions and feelings, occurring in situations close to death or in other situations of imminent physical or emotional danger (Greyson, 1983; Parnia et al., 2014). Common themes of NDE comprise, but are not restricted to, out-of-body experiences, visual and auditory hallucinations and distortion of time perception, including increased speed of thoughts (Greyson, 1983).

The neuronal mechanisms of NDE are poorly understood (Peinkhofer, Dreier & Kondziella, 2019). Nelson and colleagues previously proposed the concept that rapid eye movement (REM) sleep intrusion and REM related out-of-body experiences could occur at the time of a life-threatening event and might explain many elements of NDE (Nelson et al., 2006; Nelson, Mattingly & Schmitt, 2007). REM sleep is defined by rapid and random saccadic eye movements, loss of muscle tone, vivid dreaming, and cortical activation as revealed by desynchronization of the scalp electroencephalography (EEG). REM state features can intrude into wakefulness, both in healthy individuals and patients with narcolepsy. This may cause visual and auditory hallucinations at sleep onset (hypnagogic) or upon awakening (hypnopompic) and muscle atonia with sleep paralysis and cataplexy (Scammell, 2015). According to the hypothesis of Nelson and colleagues, danger provokes the arousal of neural pathways that, when stimulated, are known to generate REM-associated responses. This was interpreted as a “diathesis stress model” (Nelson et al., 2006; Long & Holden, 2007). In this model, an unusually sensitive arousal system (i.e. the diathesis), as evidenced by the experience of REM intrusion, would predispose people to NDE in situations of stress and danger. To test their hypothesis, Nelson and colleagues conducted a survey comparing a group of individuals with self-reported NDE and an age- and sex-matched control group (Nelson et al., 2006). The results suggested that episodes of REM intrusion are more common in individuals with NDE.

The study by Nelson et al. has been criticized (Long & Holden, 2007), however, which recently inspired us to carry out a follow-up study in a different setting to address some of the criticism (Kondziella, Dreier & Olsen, 2019). For example, Long and Holden pointed out that 40% of the people with NDE in the Nelson study denied ever having experienced an episode of REM intrusion, suggesting that there may be a link between the two phenomena, but not a 1:1 relationship (Long & Holden, 2007). In our crowdsourced survey, 106 of 1034 participants reported NDE according to a Greyson NDE Scale (GNDES) score ≥7, and 50 (47%) of these individuals fulfilled the criteria of REM intrusion according to almost the identical questionnaire that Nelson and colleagues had used (Kondziella, Dreier & Olsen, 2019). In contrast, only 17% of individuals without NDE reported REM intrusions. Based on multivariate regression analysis, we found that REM intrusion is a predictor of NDE (Kondziella, Dreier & Olsen, 2019). Thus, we confirmed the results of Nelson and colleagues, but also the limitation that this is not a 1:1 relationship.

A more central point of criticism was related to the control group in Nelson and colleagues’ study which consisted mainly of medical personnel, a likely selection bias (Long & Holden, 2007). We countered this in our survey with a crowdsourced approach in which the control group originated from the same population as the NDE group (i.e. unprimed lay people) (Kondziella, Dreier & Olsen, 2019). Our survey was announced under the headline “Survey on Near-Death Experiences and (Related Experiences)”, but we did not provide further information about the content of the study. Participants were informed that their monetary reward was fixed, regardless of whether they would report having had an NDE or not. Then, we asked the participants to complete a questionnaire comprising demographic information, followed by the questions about REM intrusion. Subsequently, participants were asked if they ever had experienced an NDE. If not, the survey ended there; if yes, participants were asked in detail about this experience and information about all 16 GNDES items was collected (Kondziella, Dreier & Olsen, 2019). In this way, we think that we were able to dispel the previous criticism regarding the control group. Long and Holden also explained how the questionnaire for REM intrusion could be misinterpreted by people with NDE, possibly leading to an overestimation of the association between REM intrusion and NDE (Long & JM, 2007). It is indeed difficult to address this problem with a questionnaire containing only closed questions. Therefore, we also gave our participants the opportunity to describe their experiences in their own words (Kondziella, Dreier & Olsen, 2019).

Another approach to address this problem is to investigate if comorbidities of REM intrusion, which might be easier to detect with a questionnaire, are associated with NDE too. In this context, it is interesting that REM sleep abnormalities have been linked to migraine. Thus, recurrent vivid dreams are associated with migraine attacks (Lippman, 1954), migraine attacks often occur during REM sleep (Levitan, 1984), and migraine patients exhibit hallucinations (Lippman, 1951, 1953; Daniel C & Donnet A, 2011), increased REM sleep and prolonged REM sleep latencies (Drake et al., 1990), and they show a significantly increased frequency of dream-enacting behavior (Suzuki et al., 2013). In addition, several studies found an association between migraine and narcolepsy, a disorder involving REM intrusion (Dahmen et al., 1999, 2003; Longstreth et al., 2007; Suzuki et al., 2015; Yang et al., 2017). For example, Yang and colleagues found a consistently higher risk of developing narcolepsy in children with migraine compared to those without, and this risk was particularly high in children with migraine with aura (Yang et al., 2017).

On this basis, we hypothesized that, analogous to an unusually sensitive arousal system underlying REM intrusion, an increased susceptibility of the brain to spreading depolarization (SD), the assumed pathophysiological correlate of migraine aura (**Figure 1A**), could predispose people to NDE. To test this hypothesis, we recruited a large global sample of laypersons and investigated if the lifetime occurrence of migraine aura is more common in people with NDE.

**Figure 1.**
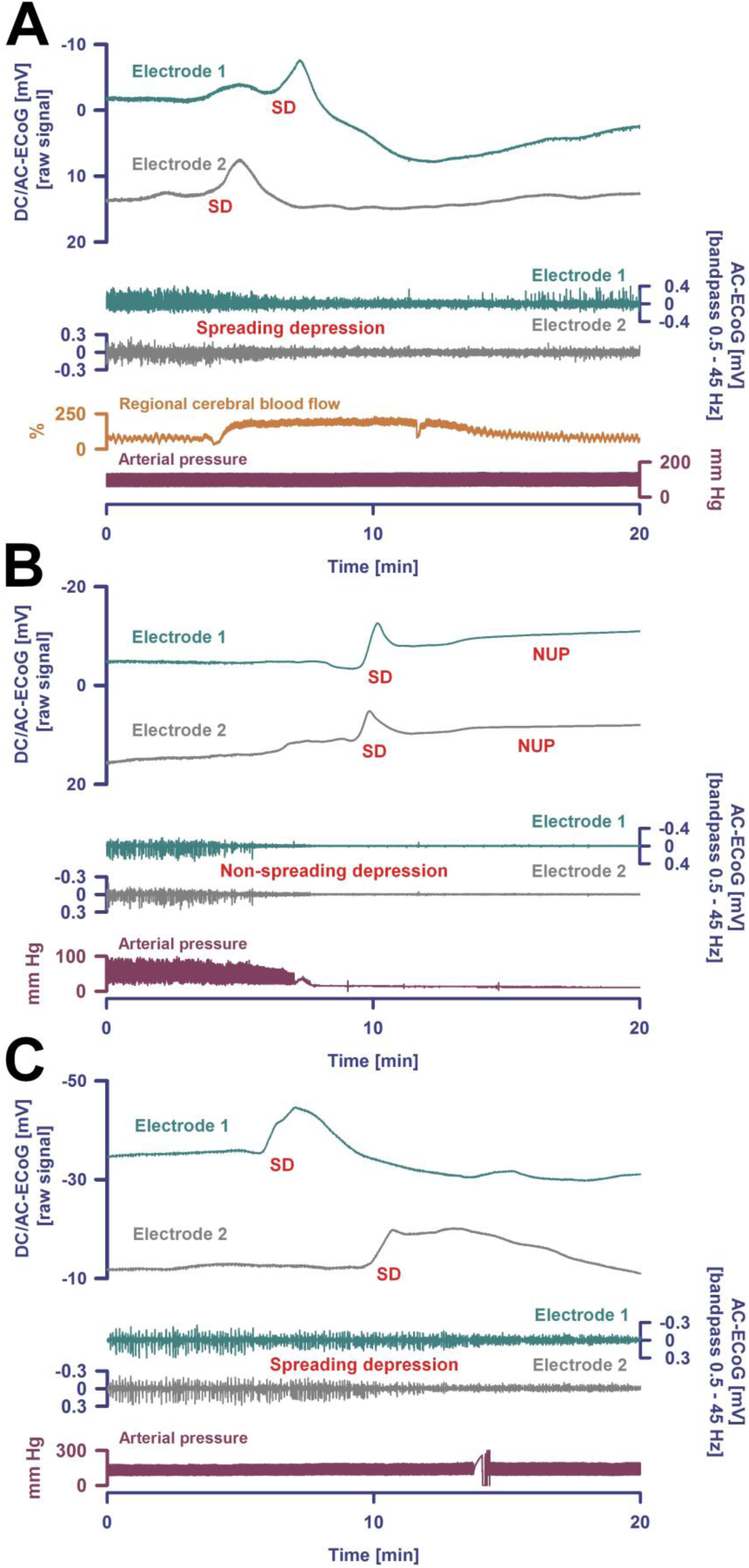
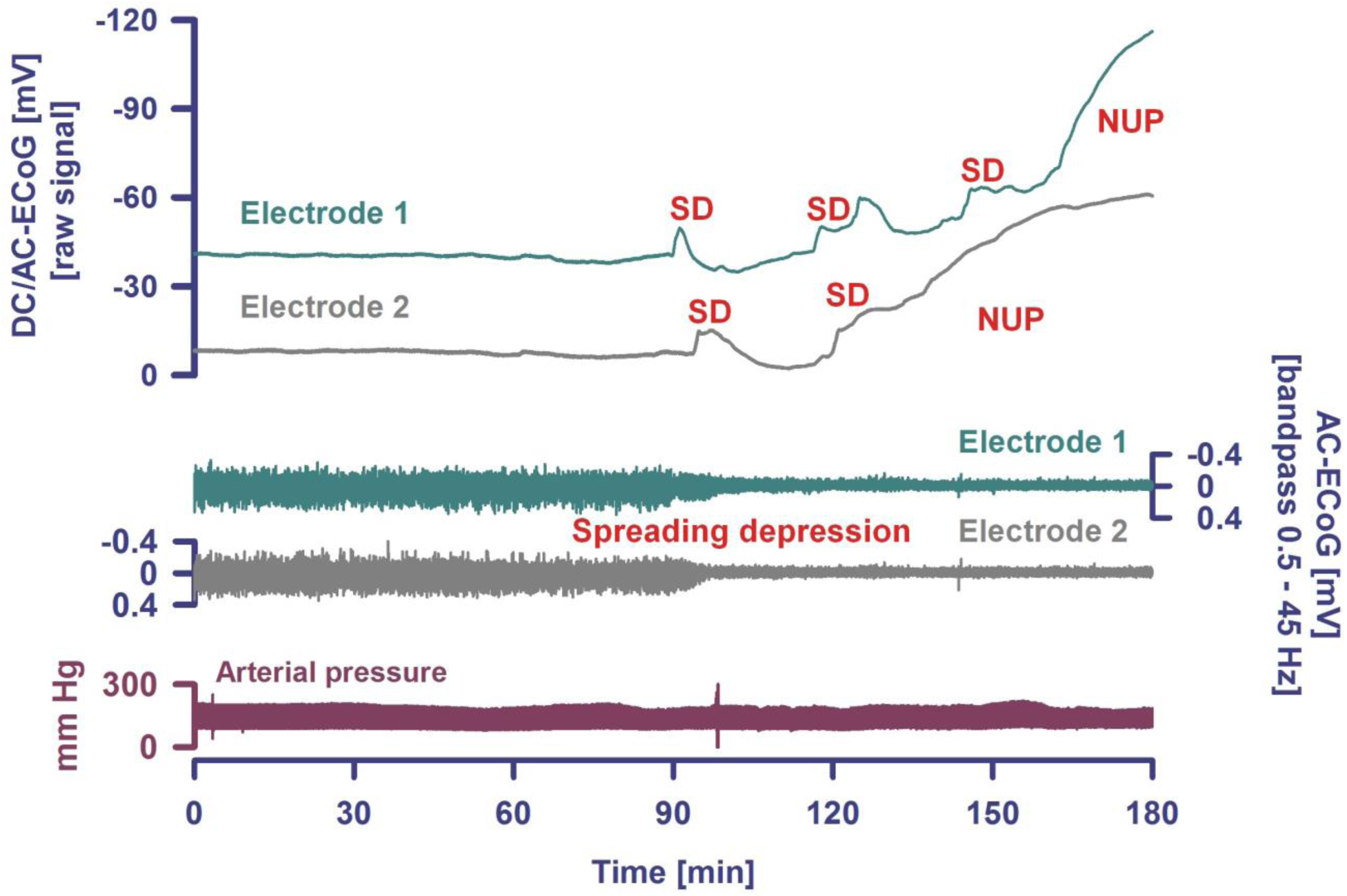
**(A)** Spreading depolarization (SD) is observed as a large negative direct current (DC) shift propagating between different electrodes (upper two traces: subdural full-band DC/alternate current (AC)-electrocorticography (ECoG) between 0 and 45 Hz, electrode separation: 1 cm) (Dreier et al., 2017). This short-lasting SD was recorded in a patient with aneurysmal subarachnoid hemorrhage (aSAH) in a metabolically largely intact and sufficiently perfused neocortex region. Based on measurements of regional cerebral blood flow (rCBF) using intracarotid ^133^ Xe and positron emission tomography, blood-oxygen-level dependent (BOLD) imaging with functional magnetic resonance imaging (MRI) and magnetoencephalography (MEG), it is assumed that the SD underlying a migraine aura should be largely similar (Dreier & Reiffurth, 2015). The patient’s perception of a migraine aura is presumably triggered by the SD-induced spreading depression of spontaneous activity (Dreier & Reiffurth, 2015), which is shown here in traces 3 and 4 as a transient reduction in amplitudes propagating between electrodes (frequency band: 0.5 - 45 Hz). It should be noted, however, that a patient can only perceive a migraine aura if this spreading depression propagates through an eloquent region of the brain (Dreier & Reiffurth, 2015). SD is characterized by the almost complete collapse of ion gradients across cell membranes, causing water influx and an almost complete loss of Gibbs’ free energy contained in the ion gradients (Dreier et al., 2013). Recovery from SD requires activation of adenosine triphosphate (ATP)-dependent membrane pumps, in particular Na,K-ATPases. Therefore, tissue ATP declines by ∼50 % during SD, not only in energy-deprived but also in well-nourished tissue (Dreier & Reiffurth, 2015). Consequently, rCBF significantly increases in normal tissue to meet the enhanced energy demand and to clear the tissue of metabolites (trace 5, measurement of rCBF using an optoelectrode and laser-Doppler flowmetry). The regional hyperemia is variably followed by a mild rCBF decrease (oligemia) during which the vascular reactivity is disturbed. The short initial hypoperfusion is an abnormality here that indicates mild impairment of the neurovascular coupling in the context of aSAH (Dreier & Reiffurth, 2015). The arterial blood pressure (trace 6) measured in the radial artery was stable during the SD. **(B)** The second patient died from hepatorenal failure several days after aSAH. (Dreier et al., 2018). Trace 5 shows the circulatory arrest, which is evidenced by the drop in arterial blood pressure. About 35 seconds after the circulatory arrest, the AC-ECoG in traces 3 and 4 begin to show the non-spreading depression of spontaneous activity (asterisk). Phase 2 lasts 95 seconds at electrode 2. Thereafter, the terminal SD occurs and spreads further from electrode 2 to electrode 1 (electrode separation: 1 cm). Terminal SD consists of the initial SD component and the late negative ultraslow potential (NUP). It remains speculative if NDE can occur in ECoG phases 1, 2 or 3. According to current knowledge, however, the occurrence of NDE in phases 2 or 3 cannot be ruled out. As explained in the main text, ECoG and scalp electroencephalography (EEG) show a flat line in phase 2, but experiments in animals and brain slices with sophisticated electrophysiological techniques including patch-clamping have shown that the synaptic terminals remain highly active in this phase and the neurons are polarized (Müller & Somjen, 2000; Fleidervish et al., 2001; Allen, Rossi & Attwell, 2004; Revah et al., 2016). Therefore, we cannot exclude with certainty that patients may experience a perception at that stage. The terminal depolarization takes place in phase 3. It cannot be excluded either that this may be associated with bright light phenomena or tunnel vision similar to what occurs during a migraine aura. Brain cells die only gradually in phase 4 which is characterized by the NUP. **(C)** After onset of the terminal cluster of SDs shown in the figure, this patient with aSAH was found to have a loss of brainstem reflexes with fixed dilated pupils, indicating the development of brain death (Dreier et al., 2019). The cluster starts here at electrode 1 and propagates to electrode 2 (traces 1 and 2). The first SD occurs in electrically active tissue and therefore causes spreading depression of the spontaneous ECoG activity which also spreads from electrode 1 to 2 (traces 3 and 4). In contrast to **(A)**, however, activity depression then persists. After the first SD, a second SD occurs, which transforms into a NUP. In contrast to **(B)**, further SDs are superimposed on the NUP, whose amplitudes become smaller and smaller. Similar to **(A)**, the arterial blood pressure (trace 5) remains stable during the SDs and the NUP. The patient was terminally extubated 20 hours later and shortly thereafter a circulatory arrest developed without further SD (Dreier et al., 2019). The cases in **(B)** (Dreier et al., 2018) and **(C)** (Dreier et al., 2019) are presented here in abbreviated form to illustrate the pivotal aspects of brain death at the tissue level. The figures were not previously published. The patients were enrolled at the Charité – Universitätsmedizin Berlin in research protocols of invasive neuromonitoring approved by the local ethics committee and written informed consent was obtained from the patients’ legally authorized representative, as described previously (Dreier et al., 2018, 2019).

## Materials & Methods

### Study design

Our objective was to investigate whether people with a history of migraine aura are more likely to have NDE, and vice versa, than people without migraine aura. We used an online platform, Prolific Academic (https://prolific.ac/), to recruit an international sample of laypeople. Like Amazon’s Mechanical Turk, Prolific Academic is a crowdsourcing online platform to recruit human subjects that can be used for research purposes (Kondziella, Dreier & Olsen, 2019; Kondziella, Cheung & Dutta, 2019) and that compares favorably in terms of data quality, including honesty and diversity of participants (Peer et al., 2017). Participants were recruited without any filters except for English language and age ≥18 years, and we excluded participants who had been enrolled in our previous study on NDE and REM intrusion (Kondziella, Dreier & Olsen, 2019). The study was announced under the headline “Survey on near-death experiences and headache” using the following text: “We wish to explore the frequency with which near-death experiences occur in the public. This should take no more than 1.5 minutes on average (a little bit longer, if you have had such an experience, and a little bit less, if you haven’t). You will be paid 0.20$ after completing the survey. Please note that we might use your anonymous answers when writing a paper.”

From all participants, we collected information about age, gender, place of residence and employment status (data provided automatically by Prolific Academic); if they had frequent headaches; if yes, if these headaches could last longer than 4 hours and were associated with visual or non-visual aura (Kaiser et al., 2019); if participants ever had an NDE; if yes, if this experience occurred in a truly life-threatening situation or in a situation that just felt so; if the experience was neutral, pleasant or unpleasant; and all participants with an NDE were asked to provide information about all 16 items of the GNDES, the most widely used standardized tool to identify, confirm and characterize NDE in research (Greyson, 1983). Like in our previous study (Kondziella, Dreier & Olsen, 2019), NDE was defined by a GNDES score ≥7. Participants with an NDE (and those who claimed an NDE but scored 6 or less points on the GNDES) were also given the opportunity to describe this in their own words (optional). **Table 1** provides details.

**Table 1.**
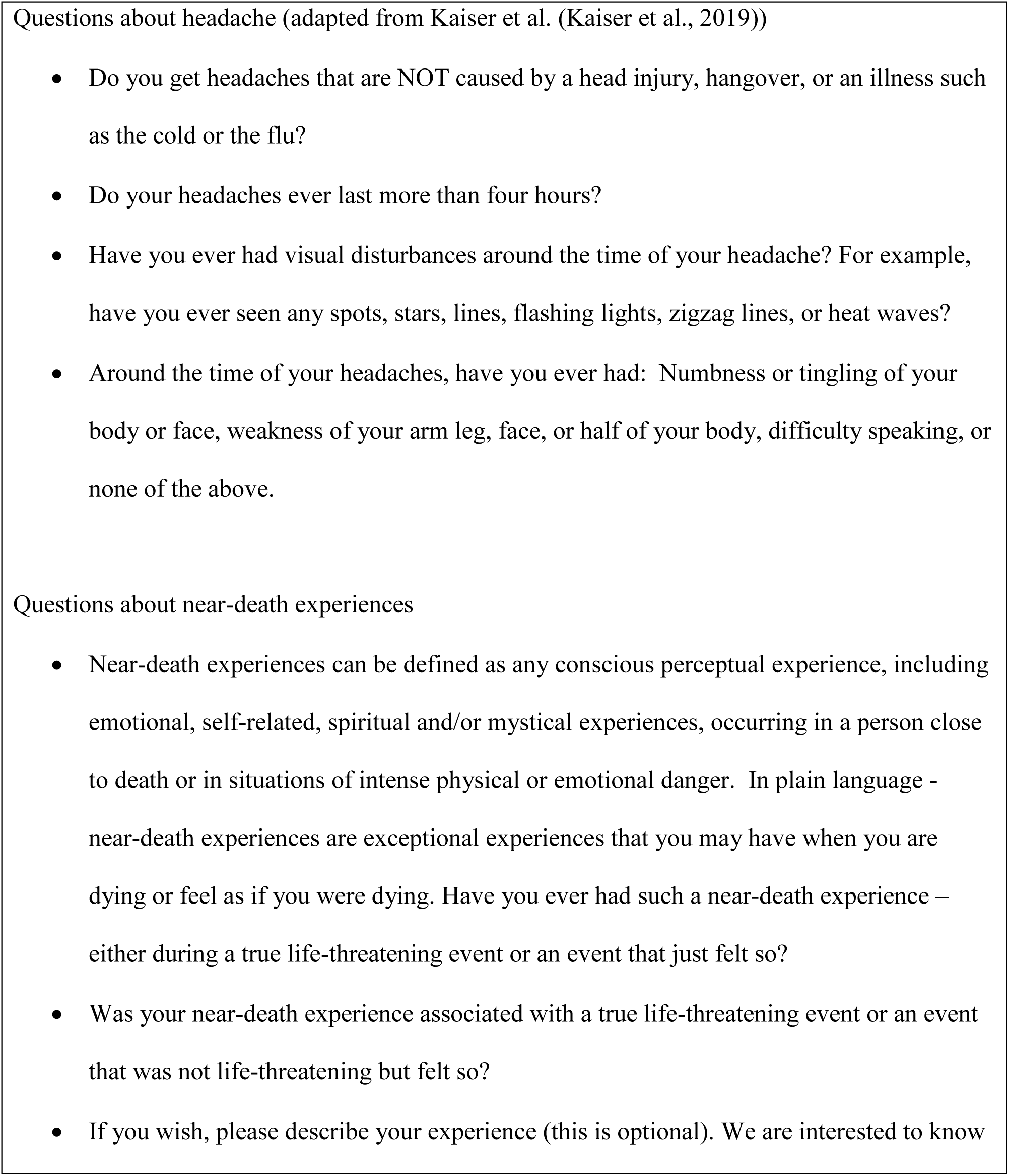
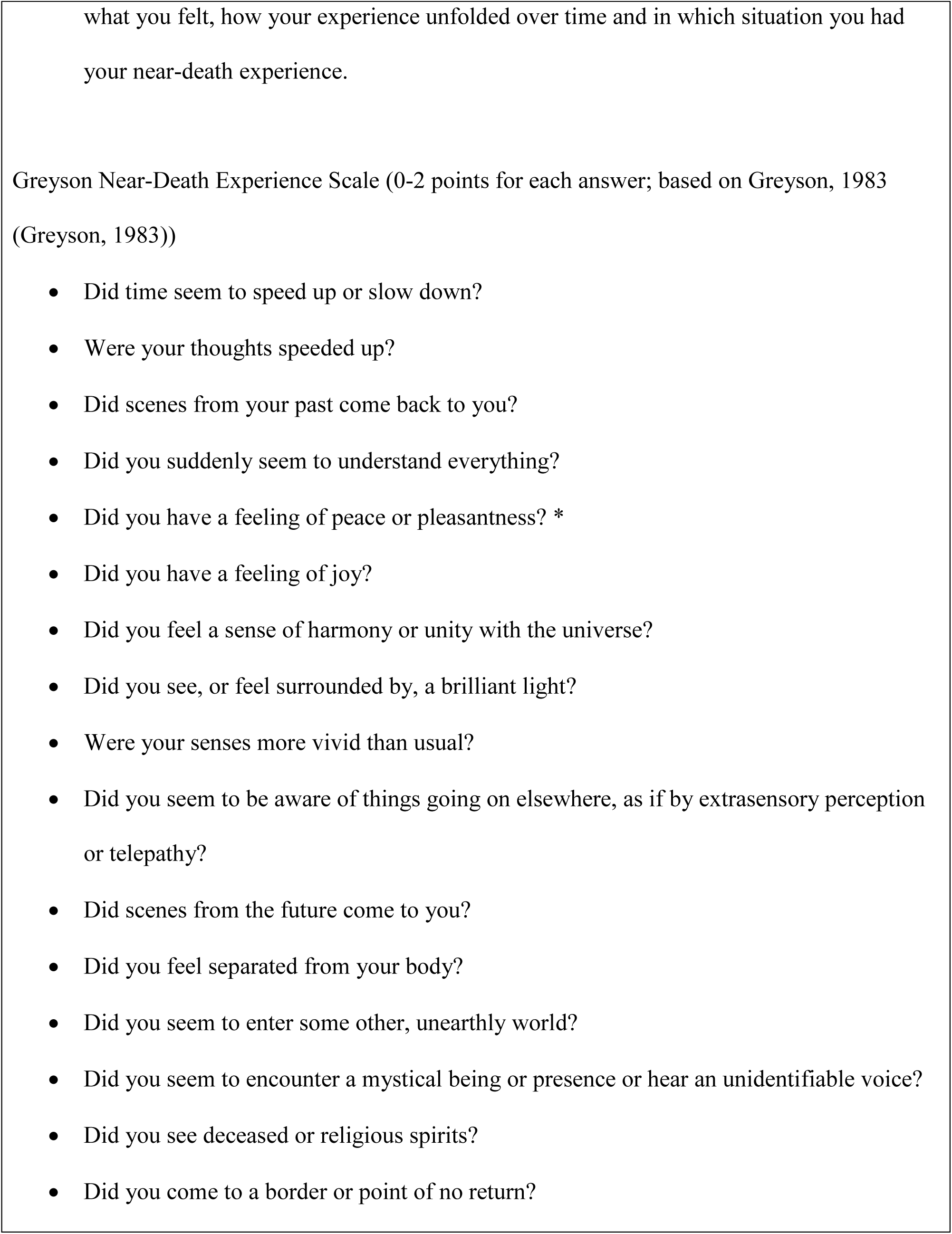
Questionnaire on headaches, migraine aura and near-death experiences. * In contrast to the Near-Death Experience Scale, we also questioned about unpleasant experiences

### Statistics

Using a very high population size (300,000,000), a confidence level of 95% and a margin of error of 5%, we estimated the required sample size to be 384 participants. However, since previous studies have estimated the frequency with which NDE occur in the public to be 5-10%, including our own on NDE and REM intrusion (Kondziella, Dreier & Olsen, 2019), we decided to enroll approximately 1000 participants to identify an estimated number of 100 individuals with an NDE.

In univariate analysis, associations between potential predictors (age, gender, migraine aura) for NDE were examined using chi-square test and t-test for independent samples. Additionally, we used multiple logistic regression to analyze the association between migraine aura and NDE adjusted for age and gender. The level of significance was 0.05 (two-sided) for all statistical tests. Statistical analysis was performed with SPSS 23.0 (IBM, Armonk, NY, USA).

### Ethics

Participants gave consent for publication of their anonymous data. Participation was voluntary, anonymous and restricted to those aged 18 years or older. Participants received a monetary reimbursement after completing the survey, in accordance with the Prolific Academic’s ethical rewards principle (≥ $6.50/h). The Ethics Committee of the Capital Region of Denmark waives approval for online surveys (Section 14 (1) of the Committee Act. 2; http://www.nvk.dk/english).

### Data Availability Statement

The de-identified raw data are provided in the *online supplemental files*.

## Results

We recruited 1037 laypeople from 35 countries (mean age: 31 years, standard deviation: 11.1 years, median age: 28 years, interquartile range (IQR): 23-36 years; 76% fully or part-time employed or in training), most of which were residing in Europe and North America (**Figure 2**). 531 participants (52%) identified themselves as female, 500 (48%) as male and six as transgender.

**Figure 2.**
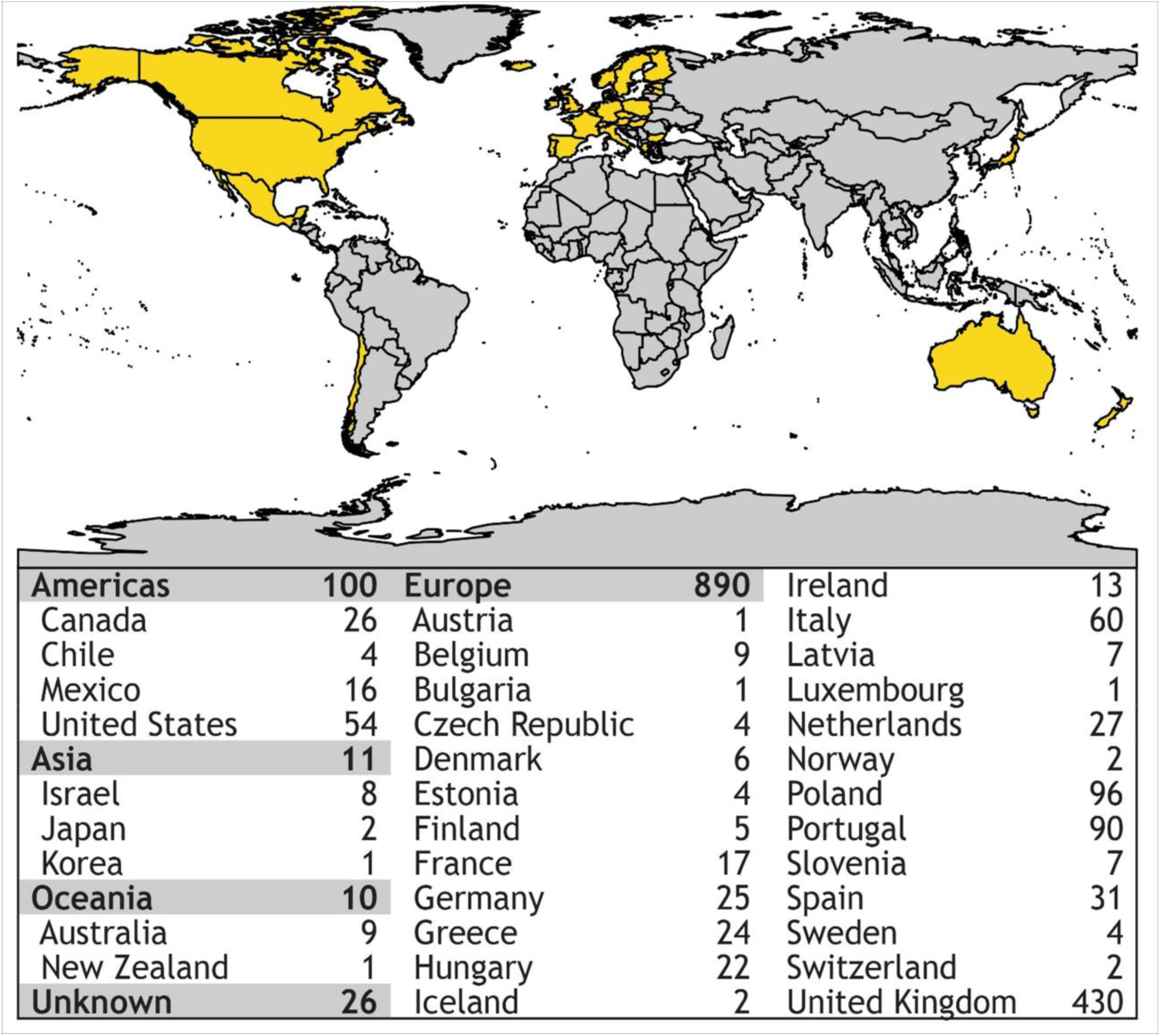
Using an online crowdsourcing platform, we recruited 1.037 lay people from 35 countries on 5 continents, the majority from Europe and North America.

### Near-death experiences: Frequency and phenomenology

Two-hundred-eighty-six participants (28%; CI 95% 25-30%) claimed an NDE. The most frequent symptoms were abnormal time perception (faster or slower than normal; reported by 257 participants; 90%); extraordinary speed of thoughts (n=169; 59%); exceptional vivid senses (n=165; 58%); and feeling separated from one’s body, including out-of-body experiences (n=113; 40%). Participants perceived the situation in which they made their experience slightly more often as truly life-threatening (n=165; 58%) than not (n=121; 42%).

However, only 81 of 286 individuals who claimed an NDE reached the threshold of ≥7 points on the GNDES (28%; CI 95% 23-34%). Hence, confirmed NDE were reported by 81 of 1037 participants (8%; CI 95% 6.3-9.7%) (**Figure 3**). Confirmed NDE were perceived much more often as pleasant (n=29; 49%) than experiences that did not qualify as NDE according to the GNDES (n=21; 13%; p<0.0001; Chi-square test; neutral experiences excluded). **Table 2** provides selected written reports from participants with an NDE of ≥7 GNDES points and **Table 3** from participants with <7 GNDES points.

**Figure 3.**
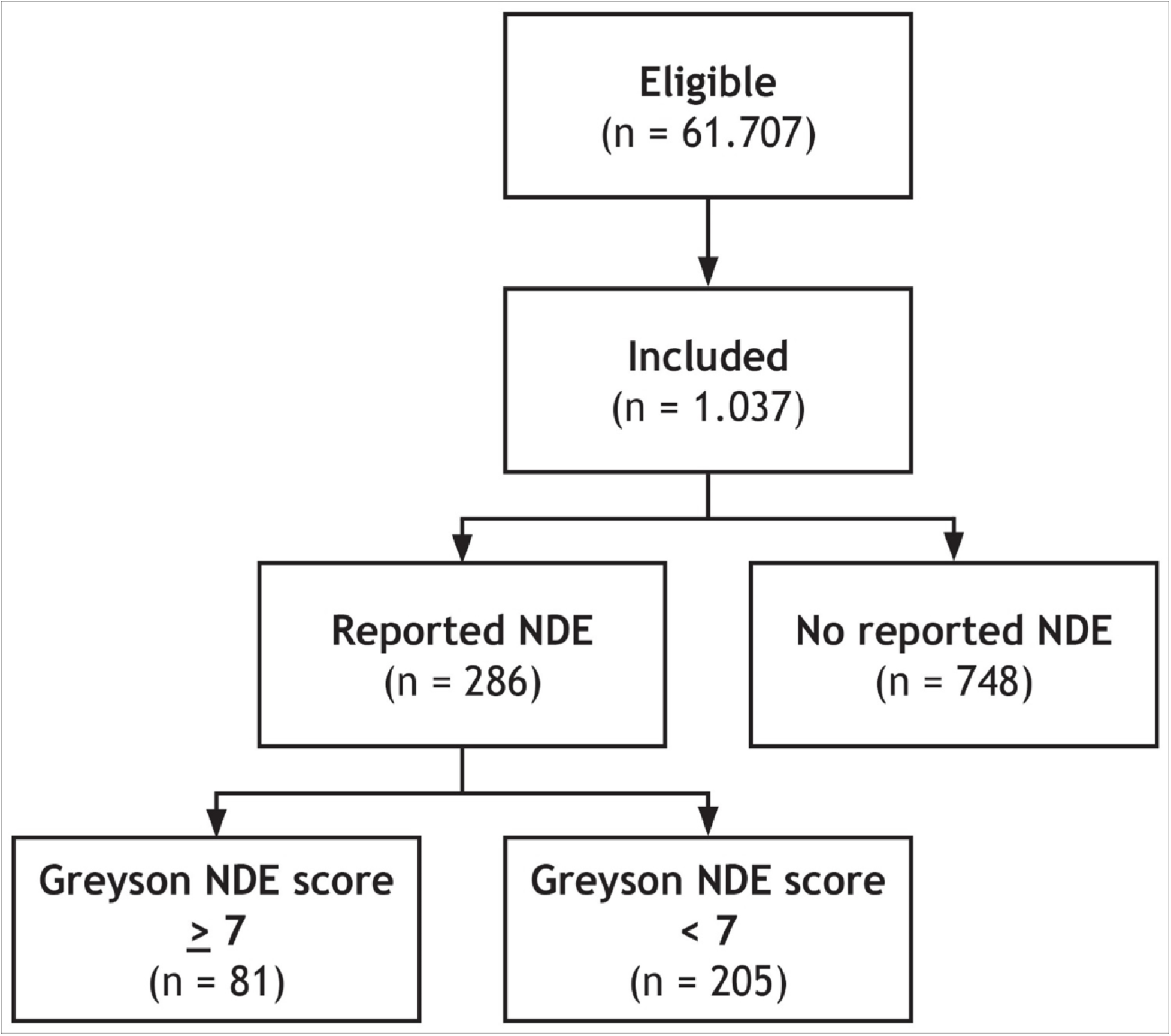
Of 61.707 eligible lay people registered with Prolific Academic (https://prolific.ac/; accessed on February 4, 2019), we enrolled 1.037 participants; 81 (7.8%; CI95% 6.3-9.7%) of whom reported a near-death experience that fulfilled established criteria (Greyson Near-Death Experience Scale score of 7 or higher). n = number of participants; NDE – near-death experience

**Table 2.**
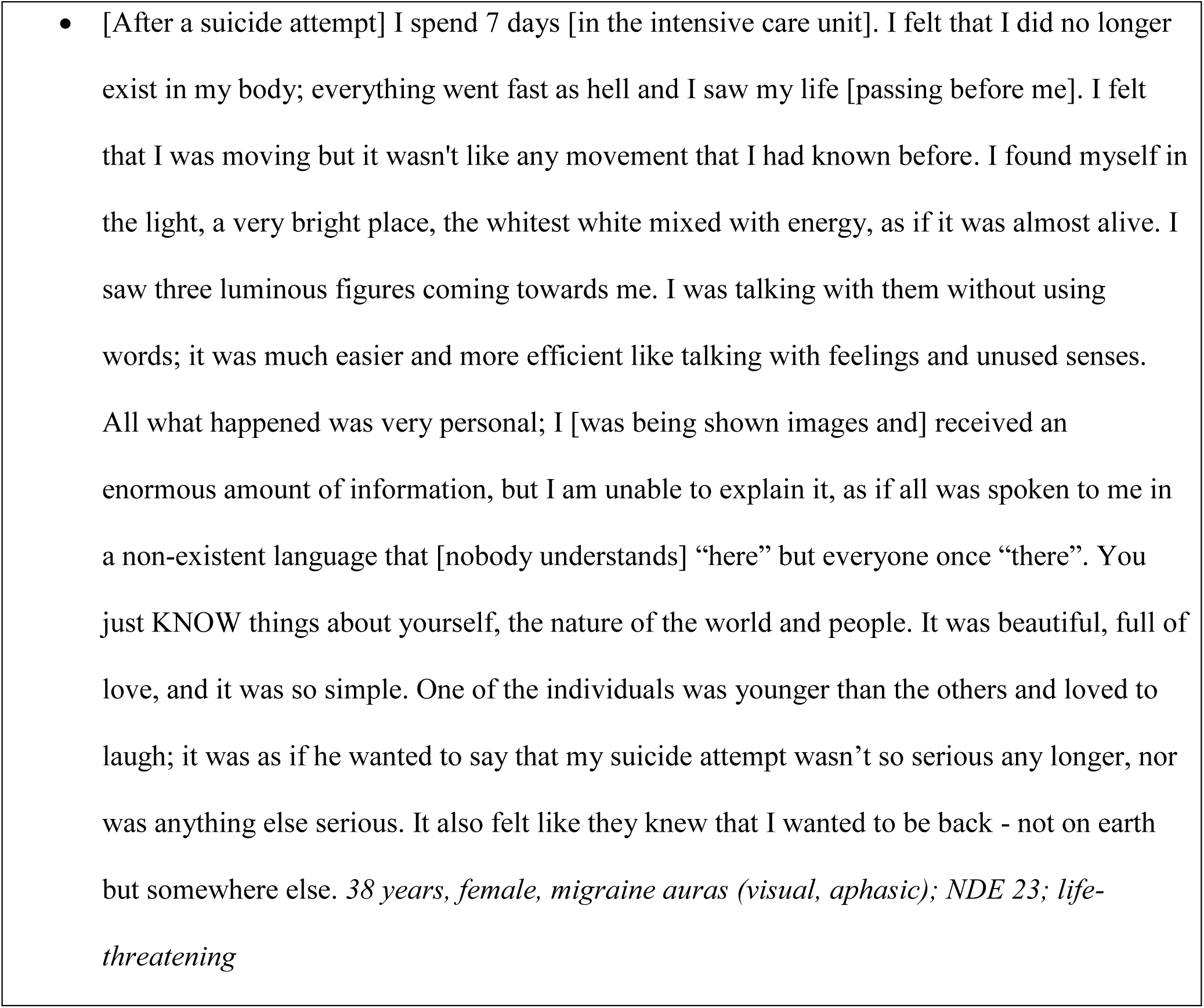
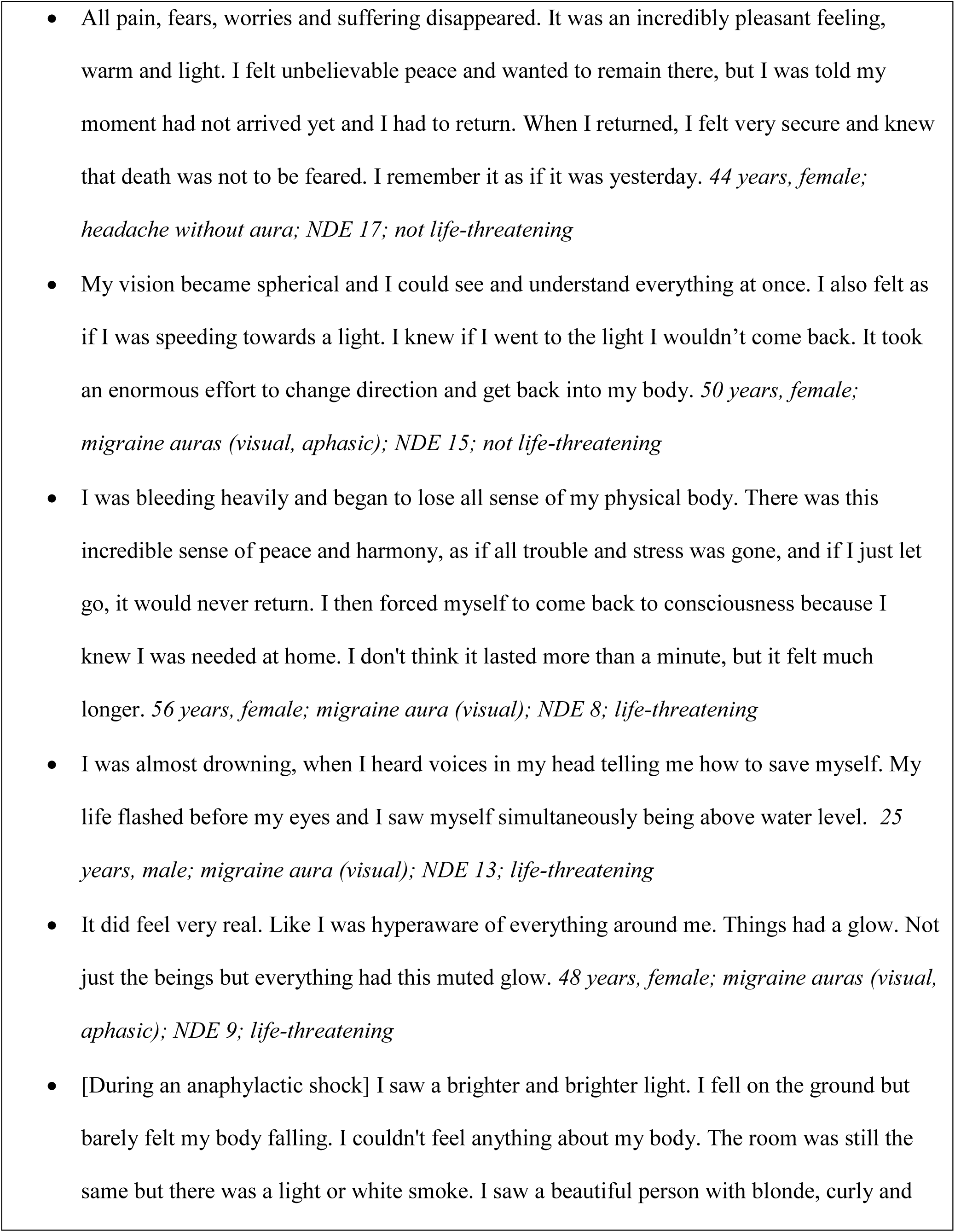
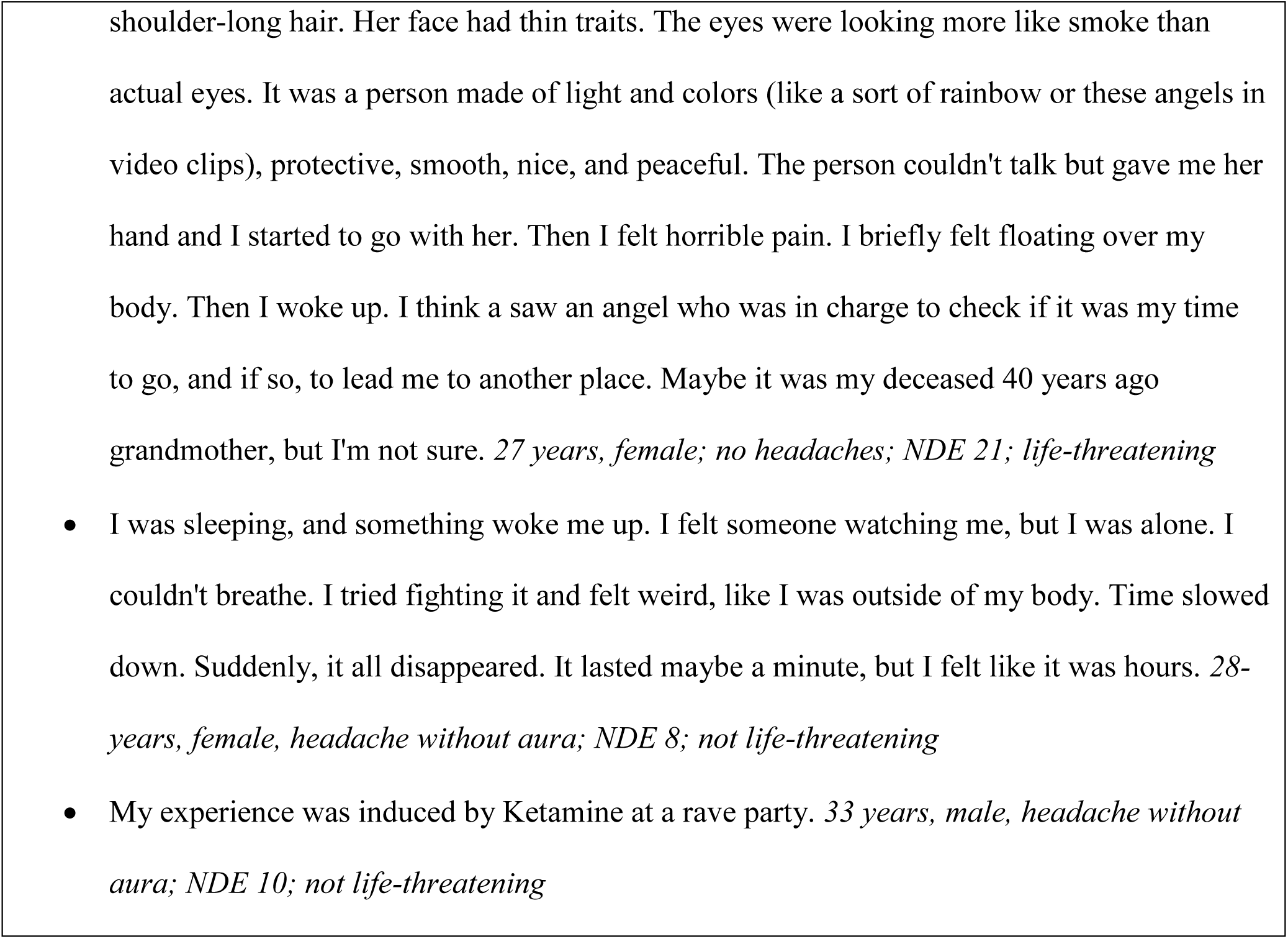
Selected reports from participants with an experience that reached the threshold of ≥7 points on the Greyson NDE scale to qualify as a near-death experience. Note that the last two comments describe experiences during ingestion of ketamine (which has been suggested as the chemical most likely to cause drug-induced near-death experiences (Martial et al., 2019)) and REM sleep disturbance (which has been identified in another recent study as a likely mechanism of near-death experiences (Kondziella, Dreier & Olsen, 2019)). Comments are edited for clarity and spelling.

**Table 3.**
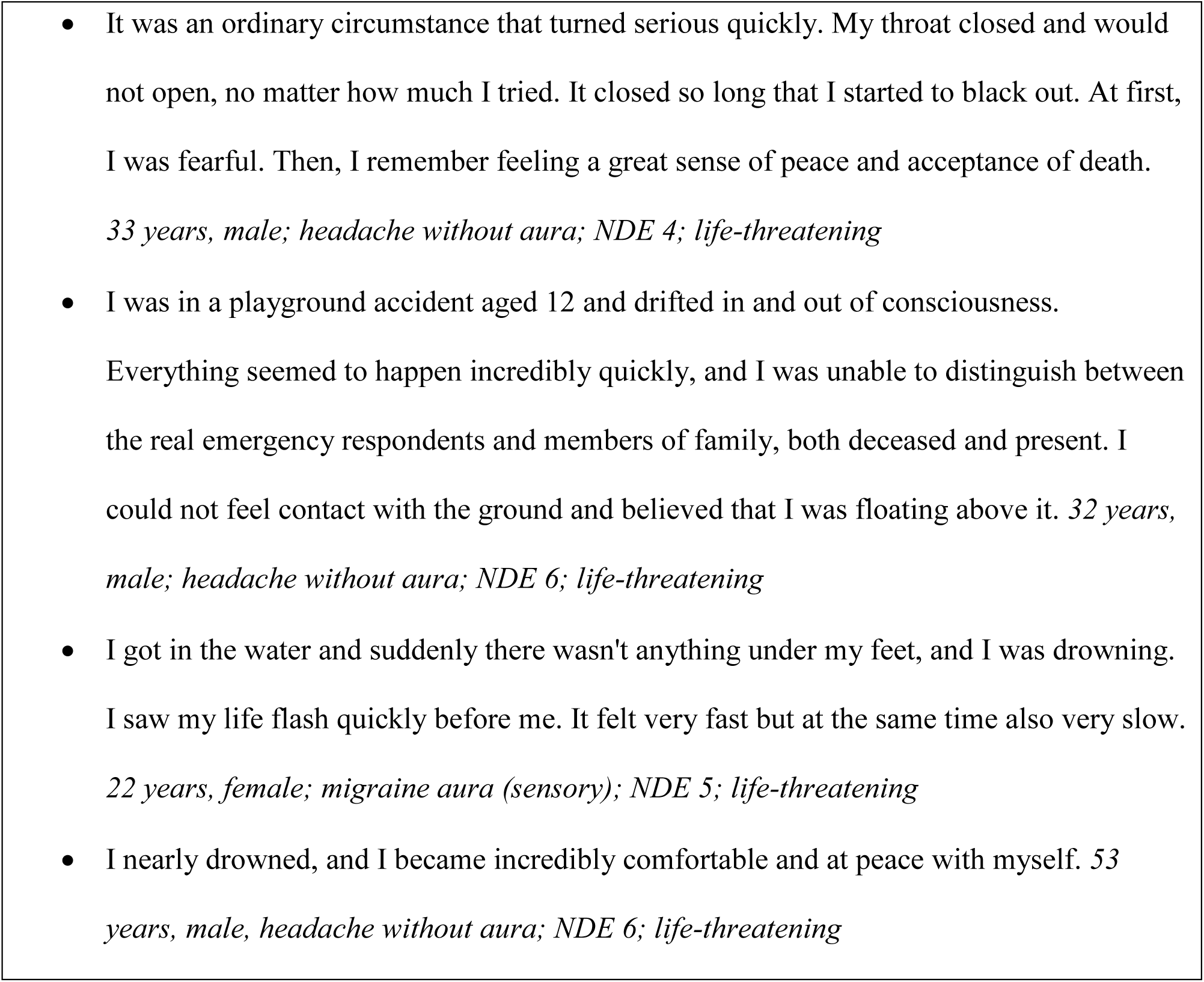
Selected reports from participants with an experience below the threshold of ≥7 points on the Greyson NDE scale. Comments are edited for clarity and spelling.

### Headache and migraine aura

Seven-hundred-twenty of 1037 individuals (69%) answered “yes” to the following question about a primary headache disorder: “Do you get headaches that are NOT caused by a head injury, hangover, or an illness such as the cold or the flu?” The male-to-female ratio of people who responded “yes” to this question was 1:1.3. Two-hundred-fifty-four of 1037 individuals (24%) fulfilled criteria (Kaiser et al., 2019) of having experienced a migraine aura at any point during their lifetime. Individuals could have different types of migraine aura. Two-hundred-thirty of 254 (91%) individuals reported having had visual auras, 60 (24%) somato-sensory auras, 49 (19%) motor auras and 21 (8%) aphasic/dysarthric auras. Hundred-seventy-four of 531 women (33%) had a migraine aura and 77 of 500 men (15%). This difference was statistically significant (p<0.001, Chi-square test; male-to-female ratio: 1:2.2). People with migraine aura were slightly older than people without (median age: 30 (IQR: 24-38) years versus median age: 28 (IQR: 22-36) years, p=0.005, Mann-Whitney Rank Sum Test).

### Near-death experiences and evidence of migraine aura

There were no significant associations between confirmed NDE and age (p>0.6, t-test independent samples) or gender (p>0.9, chi-square test). The only significant association was between confirmed NDE and migraine aura: Forty-eight (6.1%) of 783 subjects without migraine aura and 33 (13.0%) of 254 subjects with migraine aura had experienced an NDE (p<0.001, chi-square test, odds ratio = 2.29). In multiple logistic regression analysis with age, gender and the interaction of age and gender, none of these potential predictors was significant. However, migraine aura remained significant after adjustment for age (p<0.001, odds ratio = 2.31), gender (p<0.001, odds ratio = 2.33), and both age and gender (p<0.001, odds ratio = 2.33).

## Discussion

### The prevalence of NDE

The prevalence of individuals with an NDE is estimated at about 4-8% in the general population (Gallup G, 1982; Knoblauch, Schmied & Schnettler, 2001; Perera, Padmasekara & Belanti, 2005; Facco & Agrillo, 2012; Chandradasa et al., 2018). In our survey it was 8%. We found a prevalence of 10% using the same criteria in our previous crowdsourcing online survey on NDE and REM intrusion (Kondziella, Dreier & Olsen, 2019), indicating that this prevalence is quite robust. Unlike most previous reports in which NDE were almost always associated with peace and well-being (Thonnard et al., 2013; Charland-Verville et al., 2015; Martial et al., 2017, 2018; Cassol et al., 2018), we confirmed our earlier findings that many people find their NDE unpleasant (Kondziella, Dreier & Olsen, 2019). However, experiences with the cut-off score of ≥7 GNDES points were reported significantly more often as pleasant (49%) than experiences with a lower score (13%).

### Migraine aura is a predictor of NDE

Migraine aura was a predictor of NDE in our sample. This association was very stable. Regardless of whether either no adjustment, an adjustment for age, for sex or for both was performed, the odds ratios for migraine aura only varied between 2.29 and 2.33. However, a potential limitation of our study is the announcement of the internet query in which we stated that we would investigate for NDE and headache. This might have attracted more people with NDE and headache. The overall prevalence for all types of primary headache, including tension-type headache, was 69% in our survey. Tension-type headache is the most common form of headache (Jensen, 2018). Its aggregate prevalence in the general population across different studies was 38% (Jensen, 2018). Yet, in a population-based study in Denmark, a much higher lifetime prevalence of 78% was found (Lyngberg et al., 2005; Jensen, 2018). The high prevalence of primary headaches in our survey is hence within the realm of possibility but raises the question if we have attracted a disproportionate number of people with headache. This could include people with migraine with aura. The observation that 24% of the participants in our survey met criteria for a migraine aura, while population-based studies have estimated this prevalence at only 4% in the general population, renders this indeed likely (Russel et al., 1995). The young average age, typical of an Internet-based study, could have contributed to over-representation of migraineurs with aura. The way we phrased our headache questions could be another reason, as we did not intend to validate a migraine diagnosis according to established criteria (Kaiser et al., 2019). Instead, we used a more inclusive approach to identify people with a high likelihood of having migraine aura because we were not interested in migraine *per se* but rather in migraine aura as a possible predictor for an NDE (Kaiser et al., 2019). Since population-based studies suggest that spontaneous migraine aura is four times less common in people without typical migraine headache than in people with typical migraine headache (Russel et al., 1995), it is unlikely that the over-representation of people with migraine aura in our survey resulted from the fact that we also included people with migraine aura without typical migraine headache. However, we did not ask whether the aura symptoms lasted at least 5 minutes. (It should be noted that the threshold of >5 minutes to classify as migraine aura is arbitrary. Accordingly, in humans it has been shown that SD, the pathophysiological correlate of migraine aura, may occur in spatially very limited fields and that the propagation speed in the cortical tissue ranges between ∼2 and 9 mm/min (Woitzik et al., 2013)). On one hand, this could have contributed to the discrepancy between our data and population-based migraine studies. On the other hand, the male-to-female ratio in individuals with migraine aura was 1:2.2 in our survey, which is well in line with the results of population-based studies and supports that we indeed detected variants of migraine aura (Russel et al., 1995). In contrast, the male-to-female ratio of a primary headache disorder, be it tension-type headache, migraine or a rarer headache, was 1:1.3 overall. This ratio is well in line with the assumption that the vast majority of primary headache sufferers in our survey had episodic tension-type headache (Jensen, 2018).

The recurrent burden of headache may have increased motivation to participate in our survey, although this remains entirely speculative. The important question, however, is whether the combination of NDE and migraine aura disproportionately increased the motivation of affected people to join our study. Mathematically, we deal with three random factors: migraine aura (yes/no), NDE (yes/no), and participation (yes/no). The two-fold dependencies between participation and migraine aura or NDE appear unproblematic. In contrast, a three-fold dependency between participation, migraine aura and NDE could have produced a spurious association. However, we consider this unlikely because, for instance, the entire survey was finished during such a short time frame (i.e. within 3 hours after posting the survey online) that word-of-mouth communication of the survey’s topic seems very unlikely. As we cannot completely rule out this possibility, future studies will be necessary to verify that NDE and migraine aura are indeed associated. That said, Internet-based surveys and more traditional mail-based questionnaires or laboratory-based studies each have their advantages and disadvantages (Kaiser et al., 2019). We suggest that a combination of the different approaches is more meaningful than using just one method (Kondziella, Dreier & Olsen, 2019). On one side, complex clinical and ethical concepts cannot be fully captured by an online survey (Woods et al., 2015; Peer et al., 2017). On the other side, the anonymous character of a crowdsourcing online survey decreases the influence of psychological bias (Woods et al., 2015; Peer et al., 2017), because there is no incentive to satisfy the investigator by exaggerating or inventing memories. There was no monetary incentive in our survey either, since we instructed participants that their reimbursement would be the same regardless of whether they reported an NDE or headache or not. In addition, we recruited a much larger sample than would have been feasible during a conventional survey. Although participants from Europe and North America made up the largest share, ours was indeed a global sample with people from 35 countries and 5 continents.

### NDE and the neurobiology of dying

The central point in NDE research is that NDE do not only occur in healthy individuals but also during resuscitation. Thus, in the largest prospective multi-center observational trial on AWAreness during Resuscitation (AWARE), 46% of 140 survivors reported memories following their cardiac arrest with seven major cognitive themes (Parnia et al., 2014). Nine percent of the survivors met the criteria for an NDE according to the GNDES. Two percent described awareness with explicit memories of ‘seeing’ or ‘hearing’ real events related to their resuscitation. Importantly, one patient had a verifiable period of conscious awareness during which time cerebral function was not expected (Parnia et al., 2014). As surprising as this may seem, one must assume that there has to be a neurobiological basis (Nelson et al., 2006; Martial et al., 2019; Peinkhofer, Dreier & Kondziella, 2019). The pathophysiological events that occur during the process of dying are of obvious interest in this regard (Vrselja et al., 2019). The transition from life to death is thus characterized by four major events: loss of circulation, loss of respiration, loss of spontaneous electrocorticography (ECoG) activity and a terminal SD without repolarization. These four events occur always, but not necessarily in the same order (Dreier et al., 2018, 2019; Carlson et al., 2018). In the most common scenario, arrest of systemic circulation, respiration and ECoG activity develops more or less simultaneously, while terminal SD follows the complete arrest of ECoG activity with a latency of 13 to 266 seconds (Dreier et al., 2018). Along this sequence, the invasively recorded direct current (DC)/alternate (AC)-ECoG activity can be roughly divided into four different phases which are illustrated with an original recording from a previous study (Dreier et al., 2018) in **Figure 1B**: In phase 1, spontaneous ECoG activity is still measurable; phase 2 is characterized by a complete loss of ECoG activity starting simultaneously in different cortical regions and layers, which is referred to as non-spreading depression of spontaneous activity (Dreier, 2011); in phase 3, the terminal SD starts but, from a phenomenologically point of view, is initially similar to SD spreading in healthy grey brain matter (**Figure 1A**) (Dreier & Reiffurth, 2015; Hartings et al., 2017a); and finally, in phase 4 a negative ultraslow potential signals the second phase of terminal SD (Oliveira-Ferreira et al., 2010; Hartings et al., 2017b; Dreier et al., 2018, 2019; Lückl et al., 2018; Carlson et al., 2018).

The pertinent question arising from the AWARE study is whether phase 2 and (the transition to) phase 3 are compatible with a conscious perception by the patient - and hence, might contribute to the pathophysiological mechanisms of an NDE. On closer examination of the experimental data, it is interesting that the non-spreading depression of spontaneous ECoG activity in phase 2 does not result from a loss of synaptic activity, but on the contrary from vesicular release of various transmitters, including GABA and glutamate, leading to an incoherent, massive increase in miniature excitatory and inhibitory postsynaptic potentials that replace the normal postsynaptic potentials (Fleidervish et al., 2001; Allen, Rossi & Attwell, 2004; Revah et al., 2016). This probably leads to gradual depletion of the releasable pool of vesicles in the synaptic terminals, and thereby significantly distorts neuronal interactions (Fleidervish et al., 2001; Revah et al., 2016). (Not only are the miniature potentials small, but the abnormal neuronal desynchronization also prevents these potentials from summing-up, which precludes their measurement using comparatively insensitive methods such as subdural and intracortical ECoG or the even cruder scalp EEG.) Initially, neurons are hyperpolarized (Tanaka et al., 1997; Müller & Somjen, 2000). Over time, intracellular calcium and extracellular potassium concentrations gradually increase, while extracellular pH decreases (Kraig, Ferreira-Filho & Nicholson, 1983; Mutch & Hansen, 1984; Nedergaard & Hansen, 1993; Erdemli, Xu & Krnjevic, 1998; Müller & Somjen, 2000; Dreier et al., 2002). Eventually, hyperpolarization turns into neuronal depolarization. When the adenosine triphosphate (ATP) stores are exhausted, ATP-dependent membrane pumps such as the Na,K-ATPase become unable to replenish the leaking ions. Consequently, SD erupts at one or more sites of the cortical tissue and spreads into the environment as a giant wave of depolarization. It is important to understand that this terminal SD marks the onset of the toxic cellular changes that ultimately lead to death, but it is not a marker of death *per se*, since the SD is reversible – to a certain point – with restoration of the circulation (Hossmann & Sato, 1970; Heiss & Rosner, 1983; Memezawa, Smith & Siesjö, 1992; Ayad, Verity & Rubinstein, 1994; Shen et al., 2005; Pignataro, Simon & Boison, 2007; Nozari et al., 2010; Lückl et al., 2018). Thus, in contrast to what happens during coma or sedation, when the brain dies, it undergoes a massive and unstoppable depolarization process (and hence, a very last state of “activation”) (Dreier, 2011).

Returning to the association between NDE and REM intrusion, it would be interesting to know if also a link exists between miniature excitatory/inhibitory postsynaptic potentials and REM sleep. Information is scarce, but there is indeed evidence that these potentials occur in the healthy brain and are involved in the sleep-wake cycle and both REM and non-REM sleep (Yang & Brown, 2014; Christensen et al., 2014; Sangare et al., 2016). Yet, the connection between these potentials in healthy people, on one hand, and disordered neuronal processing, including NDE, on the other hand, has never been properly investigated.

Another unsolved question is if terminal SD could produce bright light phenomena and tunnel vision similar to what happens during a migraine aura, when SD spread through healthy cortical tissue. In this context, it is particularly thought-provoking that terminal SD is not always the final event, but data from so far 3 patients indicate that terminal SD can sometimes indeed precede circulatory arrest and initiate a spreading depression of spontaneous activity like that in migraineurs with aura (**Figure 1C**) (Dreier et al., 2018, 2019; Carlson et al., 2018). In contrast to migraine aura, activity then remains depressed at the time of cardiac death.

It is important to bear in mind that virtually all humans (and all animals, including insects (Spong, Dreier & Robertson, 2017)) undergo terminal SD at the end of their life, whereas only a minority of people have a migraine aura during their lifetime. Hence, although terminal SD may play a role in the development of NDE, migraine aura during lifetime is probably not required for having an NDE with a bright light at the end of life. However, people with a propensity for migraine aura may be more likely to experience terminal SD while the brain is still electrically active (**Figure 1C**). Thus, if terminal SD facilitates NDE, this would suggest that the event of a terminal SD can still be perceived and remembered.

To substantiate or dismiss these speculations, it would be necessary to fully understand how the changing polarization states of approximately 20 billion neurons in the neocortex (Mortensen et al., 2014) create the conscious awareness of an individual, an area of intense but unsolved research (Owen et al., 2006; Giacino et al., 2014; Kondziella et al., 2016; Paulson et al., 2017; Demertzi et al., 2019). This seems important because of the increasing practice of organ donation after cardio-circulatory death (DCD). In countries where DCD is practiced, physicians have reached consensus that death should occur somewhere between a few seconds and 10 minutes after loss of circulatory function (Boucek et al., 2008; Stiegler et al., 2012; Dhanani et al., 2012; van Veen et al., 2018). Thus, a survey on postmortem organ donation in the framework of the CENTER-TBI study recently revealed that as many as 10 out of 64 centers (16%) in Europe and Israel immediately begin organ retrieval from the donor after a “flat line electrocardiogram” is detected on the monitor (van Veen et al., 2018). Critical voices have been raised, however (Rady & Verheijde, 2016; Youngner & Hyun, 2019). Due to the above-mentioned uncertainties in our understanding of the dying process, we think it is indeed prudent to consider if organ removal should first be permitted when the neurons in the donor’s brain no longer exhibit synaptic transmission and alterations of their polarization state. In other words, organ harvesting should perhaps be postponed until the donor’s entire brain has unmistakably reached the negative ultraslow potential phase of terminal SD. It follows that a better understanding of NDE may be relevant to protect the interests of potential organ donors in the context of DCD.

## Conclusions and future directions

In a large global sample of unprimed laypeople, migraine aura was significantly associated with NDE, even after multivariate adjustment. The connection between migraine aura, REM intrusion and NDE is complex. For instance, the brainstem plays an important role in REM intrusion, and dream-like hallucinations such as those in REM sleep are known from people with lesions near the mesopontine paramedian reticular formation and the midbrain cerebral peduncles (i.e. peduncular hallucinations) (Galetta & Prasad, 2017), suggesting that dysfunction of the REM-inhibiting serotonergic dorsal raphe nuclei and the noradrenergic locus coeruleus facilitates REM intrusion (Hobson, McCarley & Wyzinski, 1975; Manford & Andermann, 1998; Kayama & Koyama, 2003; de Lecea, Carter & Adamantidis, 2012). A large body of evidence further indicates that the brainstem also plays an important role in the pathogenesis of migraine (Akerman, Holland & Goadsby, 2011); REM sleep abnormalities have been described in migraineurs; and several reports have substantiated the notion that migraine, in particular migraine with aura, is associated with narcolepsy (Lippman, 1951; Levitan, 1984; Drake et al., 1990; Dahmen et al., 1999, 2003; Longstreth et al., 2007; Suzuki et al., 2013, 2015; Yang et al., 2017). Hence, we and others have suggested that REM intrusion is a predictor of NDE (Nelson et al., 2006; Kondziella, Dreier & Olsen, 2019). In the present study we found that migraine aura is also a predictor of NDE. The relationship between NDE and migraine aura raises many novel questions which deserve further investigations. In the broadest sense, excitation/inhibition imbalance across different brain structures is likely to play a role (van den Maagdenberg et al., 2004; Tottene et al., 2009; Ambrosini et al., 2016). However, migraine aura also has an important vascular component that is particularly interesting for the study of NDE and the dying brain and further increases the complexity of these phenomena and their interactions (van den Maagdenberg et al., 2004; Tottene et al., 2009; Dreier & Reiffurth, 2015).

## Acknowledgements

N/A.

